# Functional Characterization of Transcriptome-Wide Isoform Switching in Hürthle Cell Carcinoma (HCC)

**DOI:** 10.64898/2026.07.23.740299

**Authors:** Razmia Sabahat Butt, Afreenish Amir, Rehan Zafar Paracha

**Affiliations:** School of Interdisciplinary Engineering & Sciences (SINES), National Univsersity of Sciences and Technology (NUST), Islamabad, Pakistan; Public Health Laboratories Division, National Institutes of Health (NIH), Islamabad, Pakistan; Faculty of Pharmaceutical Sciences (FPS), Riphah International University, Islamabad, Pakistan

## Abstract

Hürthle cell carcinoma (HCC) is an aggressive form of thyroid cancer. While mitochondrial DNA mutations and chromosomal losses have been identified in HCC, isoform switching, and its functional consequences remain uncharacterized. This study reanalyzed NCBI GEO dataset GSE228870 (n = 32), using Salmon and *IsoformSwitchAnalyzeR()* to identify isoform switching. The analysis resulted in 371 switches across 335 genes showing functional consequences including loss of protein domains, shorter open reading frames (ORFs), loss of signal peptides and novel sub-cellular localizations. Most significant isoform switches (*q*-value < 0.05, |dIF| > 0.1) were observed in *LAMA2*, *LSP1*, *MAD2L2*, *FBLN2* and *CXCL12*, implicating extracellular matrix dysregulation, DNA damage response, immune signaling and cytoskeleton regulation. These genes are expressed in normal thyroid (median TPM 20.69, 11.66, 14.79, 134.1 & 80.76). However, specific isoforms of *LAMA2* and *MAD2L2* are not expressed in normal thyroid, explaining tumor-specific expression in HCC. Alternative transcription termination site (ATTS) gain was significant, suggesting altered 3’ end in HCC transcripts. TCGA SpliceSeq showed *LSP1*, *FBLN2* and *CXCL12* undergo alternative promoter (*LSP1* exon1 PSI=94.5%, *FBLN2* exon2 PSI=99.0%) and alternative termination (*CXCL12* exon3.3 PSI=53.9%) in thyroid cancer, suggesting ATTS and alternative transcription start site (ATSS) as shared splicing dysregulation mechanisms. This is the first systematic characterization of isoform-level dysregulation in HCC.

## Introduction

Thyroid cancer is the most common endocrinological cancer, ranked as seventh most prevalent cancer globally, with incidence increasing by 200% from 1992 to 2018 (Alagoz et al. 2024). The risk factors for thyroid cancer include gene mutations, pre-existing thyroid diseases, exposure to high levels of radiation, obesity, smoking, heavy alcohol consumption, lack of exercise and poor diet (Forma et al. 2025). Thyroid cancer is classified into four main types including papillary thyroid cancer (PTC), follicular thyroid cancer (FTC), medullary thyroid cancer (MTC) and anaplastic thyroid cancer (ATC). Among these subtypes, PTC is the most common, accounting for approximately 85-90% of all thyroid cancer cases (Zhang and Xu 2024). FTC is the second most common thyroid malignancy and accounts for about 10-15% of all thyroid cancer cases (Luvhengo et al. 2023). FTC is further subdivided into three subtypes including minimally invasive, encapsulated angioinvasive and widely invasive variants. Previously, Hürthle cell carcinoma (HCC; also known as oncocytic carcinoma (OTC)) was identified as a subtype of FTC but World Health Organization (WHO) classified it as a different type of thyroid tumor (before 2017) derived from thyroid follicles (Basolo et al. 2023).

While PTC has better prognosis with overall 10-year survival rate exceeding 95% (Zhao et al. 2025), HCC is aggressive form of thyroid cancer, have greater potential for malignancy and accounts for 3-4% of thyroid cancer cases with higher occurrence in females than males (Rehman and Dhatariya 2019; Kure and Ohashi 2020). HCC is distinguished by the presence of large Hürthle cells. Hürthle cells are characterized by the presence of large eosinophilic cells with granular cytoplasm and abundant mitochondria (Sulaiman et al. 2024). The primary mode of HCC diagnosis is fine-needle aspiration cytology (FNAC). However, due to the overlapping features of HCC with other thyroid malignancies, definite diagnosis is challenging (Kim et al. 2022). There is no consensus on the treatment of HCC. Surgical removal of tumors including total or unilateral thyroidectomy is recommended for patients with 1–4cm thyroid lesions. Radioactive iodine (RAI) for treatment of HCC is still debated among researchers and general guidelines for the use of RAI in HCC are inconsistent (Kure and Ohashi 2020). Previous studies show that HCC is characterized by mutations of mitochondrial DNA (mtDNA) and chromosomal loses. The mtDNA mutations lead to impaired oxidative phosphorylation. The most common loss of function and missense mtDNA mutations occur in mitochondrial complex I-related genes such as *GRIM19/NDUFA13* and are documented in 60% of HCC cases (Singh et al. 2021; Li et al. 2025). Mutations in rat sarcoma (*RAS*) gene are observed in 15-25% cases of HCC (Singh et al. 2021). However, the main molecular effectors activated due to these genetic alterations have not been well characterized (Gopal et al. 2023).

RNA splicing is a major event in the expression of genes. Splicing involves removal of introns from pre-messenger RNA (pre-mRNA) to produce mature mRNAs. Splicing also influences other cellular mechanisms such as mRNA translation, nuclear export and can introduce premature stop codons (Bradley and Anczuków 2023). Alternative splicing (AS) results in the production of different transcripts or isoforms in humans and other higher vertebrates. Approximately 95% of multi exon genes undergo alternative splicing (Vitting-Seerup and Sandelin 2017). There are seven different types of alternative splicing events that can generate differential isoforms, including exon skipping (ES), intron retention (IR), alternative 5’ splicing (A5), alternative 3’ splicing (A3), alternative polyadenylation (AP; alternative transcription termination site (ATTS)), alternative promoters (alternative transcription start site (ATSS)) and mutually exclusive exons (MEE) (Lv et al. 2025). Isoform switching or differential isoform usage (|dIF|) refers to the production of different functional isoforms. Isoform switching is implicated in many diseases including cancer (Vitting-Seerup and Sandelin 2017). Mutations in 3’ or 5’ splice site, cis-regulatory elements and changes in gene expression can lead to production of functionally aberrant isoforms in cancer. Unlike differential gene expression analysis, isoform switching resolves the transcript level variants driving biological changes in diseases. Isoform switches have been characterized in 12 different types of cancer, including breast cancer, lung squamous cell carcinoma and thyroid cancer (Karakulak et al. 2021; Dolgalev and Poverennaya 2023). However, the data regarding isoform switching in HCC is limited and no study has documented the functional consequence of such switching.

To address the isoform-level aberrations in HCC, we reanalyzed RNA-seq dataset from NCBI Gene Expression Omnibus (GEO) (Edgar 2002) with accession ID GSE228870. The original study identified the molecular effectors activated by complex I mitochondrial mutations and chromosomal losses that are characteristic features of HCC (Gopal et al. 2023). Recent studies show that connection between cellular metabolism and alternative splicing is bidirectional. Alternative splicing can regulate metabolic pathways by generating enzymatic isoforms. Inversely, metabolic dysregulation can influence RNA processing through spliceosome activity (Voss et al. 2026). This suggests that mitochondrial dysfunction in HCC may promote transcriptional dysregulation at isoform level. Despite mitochondrial and chromosomal aberrations documented in HCC, no previous study has identified the transcriptomic dysregulation in HCC. This study uses a patient-paired design to reanalyze the dataset, identifies major isoform switches within HCC and characterizes the functional consequences of these switches, including coding potential, non-sense mediated decay (NMD) status, protein domain gain/loss, intrinsically disordered regions (IDR), sub-cellular localization and transmembrane topology. Moreover, RNA splicing events such as AS, ATTS, and ATSS usage were characterized to identify the dominant mechanisms driving isoform diversity in HCC.

## Methodology

### Dataset Selection

The secondary dataset for isoform switch analysis was obtained from NCBI GEO with accession ID GSE228870 (BioProject ID PRJNA951766). The original dataset consisted of 42 samples (18 controls and 24 HCC). However, for this analysis, paired-patient design was utilized, analyzing matched tumor and control samples. This paired-patient design led to the selection of 36 samples (18 HCC and 18 matched controls). Six HCC samples were excluded as these did not have matched controls. While 32 samples for isoform switch analysis seem limited, the paired design used here maximizes statistical power by controlling for inter-patient transcriptomic variability. Patient metadata available from NCBI GEO was limited to sample condition (tumor vs control) and patientID. No additional variables such as age, sex, tumor stage or size were available for this dataset. The complete list of samples and metadata information can be found in Supplementary Table S1. Paired-end FASTQ files sequenced on Illumina HiSeq 2000 platform, with sequence length 101 bp (average spot length 201) were obtained for selected 36 samples using fasterq-dump from SRA toolkit (Leinonen et al. 2010).

After obtaining FASTQ files, initial QC was performed using FastQC (version 0.12.1) (Andrews 2010). Quality metrics were evaluated per FASTQC’s standard HTML report. After initial QC, all-in-one processing was performed using fastp (version 0.23.4) (Chen et al. 2018) with default parameters, including automatic adapter detection, a minimum Phred quality threshold of ≥15 and a minimum post-trimming read length filter of 15bp. While the 15 bp minimum length threshold is permissive, reads shorter than the Salmon index k-mer size of 31 bp are inherently unable to seed a valid quasi-mapping and are automatically discarded during quantification, ensuring that excessively short reads do not contribute to transcript abundance estimates. The trimmed reads were obtained and used for transcript quantification.

### Transcript Quantification

Isoform-level analysis required quantification on transcript level. Transcript quantification was performed using Salmon (version 1.10.3) (Patro et al. 2017). Salmon’s quasi-mapping approach directly quantifies transcript expression against a reference transcriptome. In this step, processed FASTQ files were mapped against a reference transcriptome (human Hg38). Gencode v44 (GRCh38.p14) was selected for its comprehensive transcript coverage and compatibility with Hg38 genome assembly. The reference genome FASTA, annotation GTF and transcript files were downloaded from Gencode. The reference transcriptome was first indexed using Salmon index with k-mers of length 31. The default k-mer size suggested by Salmon for read length > 75 bp is 31. As our read length was 101bp, the default k-mer (i.e. 31) size was selected.

The index built was used to perform decoy-aware quantification. The complete reference genome was used to build decoy. Gentrome (i.e. combination of transcriptome and genome) was built and used for decoy-aware quantification. Decoy-aware aligner minimizes false mapping of reads arising from unannotated genomic regions. Validate mapping flag was enabled to enhance alignment-based verification of mappings and improve the accuracy of quantification. The library type was automatically determined by Salmon using −l A flag and later confirmed to be Inward-Stranded-Reverse (ISR) in output logs. GC bias correction (--gcBias) and sequence specific bias correction (--seqBias) were enabled to account for systematic biases inherent to Illumina sequencing. After successful mapping and quantification, mapping rate for each sample was analyzed. An additional quality filtering layer was applied at this stage and samples with mapping rate ≤ 75% were dropped. No minimum threshold was applied for million mapped reads or transcript count as importIsoformExpression() (Soneson et al. 2016; Robinson and Oshlack 2010) function of *IsoformSwitchAnalyzeR()* incorporates bias correction and performs inter-library normalization prior to isoform usage testing. To account for variations in sequencing depth and transcript length across samples, counts were normalized to transcripts per million (TPM).

### Isoform Switching Analysis

After transcript mapping and quantification, alternative isoform switch analysis and functional consequences were analyzed using Bioconductor package (version 3.20) *IsoformSwitchAnalyzeR()* (v2.6.0) in RStudio (R 4.4.1 and RStudio 2025.09.2 build 418) (Posit team 2025). Transcript abundance estimate and counts calculated by Salmon were imported into R Studio, using importRdata() function (Soneson et al. 2016) along with Gencode v44 annotation and FASTA files used for quantification. A multi factor design matrix (° condition + patientID) specifying biological condition (control vs tumor) and patient specific metadata (patientID according to the metadata information from GEO) as variables of interest was created.

After data import, pre-filtering was performed to remove single isoform genes and genes lacking consequence potential. Gene expression cutoff was set to 1 TPM and isoform expression cutoff was set to 0 (default). Principal component analysis (PCA) was performed on log_2_ transformed TPM values of the top 25% most variable isoforms to evaluate sample clustering and analyze baseline expressions using boxplots. Differential isoform usage was identified using isoformSwitchTestDEXSeq() function (Vitting-Seerup and Sandelin 2017; Ritchie et al. 2015; Anders et al. 2012) of the package on default parameters. isoformSwitchTestDEXSeq() applied negative binomial generalized linear models (GLM) to detected differential exon usage. As patientID was included in the design matrix, the function automatically accounted for individual patient confounding effects to control for inter-patient baseline variations.

Only known isoforms and coding DNA sequences (CDS) from GTF were used to identify the switches. Nucleotide and amino acid sequences of these isoforms were extracted to run the downstream analysis. Coding potential of isoforms were predicted using CPC2 (CPC 2.0 Batch Version, using genome assembly Hg38)(Kang et al. 2017), signal peptides were predicted using SignalP (SignalP 6.0, eukaryotic mode, short output no figures, fast mode) (Teufel et al. 2022), IUPred2A was used to predict IDR (on default long disorder setting) (Mészáros et al. 2018), proteins domains were predicted using Pfam (PfamScan from Galaxy.eu (Afgan et al. 2018), version 1.6, Pfam version 36.0, HMMER version 3.3.2) (Punta et al. 2011), sub-cellular localization of proteins were predicted using DeepLoc2 (version 2.0, using short output no figures and high quality model) (Thumuluri et al. 2022) and DeepTMHMM (version 1.0.57) (Hallgren et al. 2022) was used to predict protein topology. The data from these sources was imported back into R studio using their specific functions such as analyzeCPC2(), analyzeSignalP(), analyzeIUPred2A(), analyzePFAM(), analyzeDeepLoc2() and analyzeDeepTMHMM(). Alternative splicing events including intron retention, alternative transcription start, and termination sites were identified using analyzeAlternativeSplicing() function (Vitting-Seerup et al. 2014; Vitting-Seerup and Sandelin 2017) of the package.

Functional consequences of isoform switch were calculated using analyzeSwitchConsequences() function (Vitting-Seerup and Sandelin 2017) of package on default parameters to display features stemming from statistically significant isoform switches (with *q*-value < 0.05 & |dIF| > 0.1). Biologically significant consequences were selected, including intron retention, coding potential, NMD status, domain isotype, domains identified, ORF sequence similarity, IDR type, IDR identified, signal peptide identified, sub-cellular location and isoform topology. *IsoformSwitchAnalyzeR()* calculates isoform usage via isoform fraction (IF). IF represents the relative abundance of a specific isoform originating from a single gene. Mathematically, IF is calculated as:

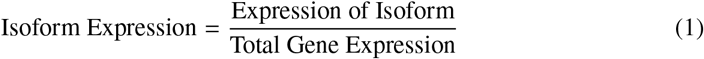

The denominator in IF represents the total expression of all isoforms of a particular gene. Consequently, IF is used to measure differential isoform fraction (|dIF|) between two conditions. |dIF| quantifies the difference in expression of isoform between two biological conditions. Mathematically, |dIF| is calculated as:

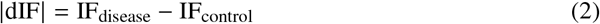

Multiple testing correction was performed using the Benjamini-Hochberg method to control the false discovery rate (FDR), with the resulting adjusted *p*-values reported as *q*-values. An isoform switch was considered significant if it met both threshold of *q*-value < 0.05 and an absolute |*dIF*| > 0.1. A |*dIF*| > 0.1 means that usage of a particular transcript has increased or decreased by 10% in disease as compared to controls. A list of most significant switches was obtained. The results of all these analyses were plotted for better visualization (Vitting-Seerup and Sandelin 2017).

The genes showing isoform switching with functional consequences in HCC were used to perform functional enrichment analysis using g:Profiler to determine their involvement in biological processes and pathways (Kolberg et al. 2023). The gene list was used to query g:GOSt with *Homo sapiens* as target organism, and Benjamini-Hochberg FDR correction method with *q*-value < 0.05 as significant threshold was applied. Gene ontology (GO) biological processes, cellular components and molecular functions were assessed along with pathway databases including KEGG, Reactome and WikiPathways.

The resulting genes and isoforms were validated using Genotype-Tissue Expression Project (GTEx) portal (Lonsdale et al. 2013) and Cancer Genome Atlas Sequence Database for Alternative mRNA Splicing (TCGA SpliceSeq) (Ryan et al. 2015). GTEx portal was used to validate the expression of genes and their respective isoforms in normal thyroid tissue. TCGA SpliceSeq was used to validate the splicing mechanisms in thyroid cancer. Top 10 genes identified from isoform switch analysis were queried in GTEx portal and TCGA SpliceSeq. The TPM values of genes and their respective isoforms from GTEx were analyzed to validate *IsoformSwitchAnalyzeR()* results. Percent Spliced In (PSI) values from SpliceSeq were evaluated for each splicing event in isoforms of target genes. Both GTEx and SpliceSeq do not harbor specific data for HCC, hence thyroid tissue and thyroid cancer data was used to validate and compare our results.

## Results

### Salmon Quant Results

The original dataset (GSE228870) consisted of 42 samples. For this reanalysis, a patient-paired design was chosen, and 36 samples were selected. Six HCC samples were dropped as they did not have matched controls. After Salmon quantification, mapping rate, total mapped reads and number of detected transcripts were analyzed. The overall mapping rate of samples ranged from 85.62% to 93.71%. Two samples showed mapping rate < 75% (SRR24053377 & SRR24053395). These tumor samples along with their matched controls (SRR24053378 & SRR24053396 respectively) were dropped to preserve paired design integrity (Supplementary Table S2), leaving 16 pairs (32 samples) for isoform switch analysis. Figure 1 shows the combined QC metrics of all samples, illustrating that majority of samples passed the mapping rate threshold of 75%, with mapped reads and transcript count exceeding 20.4M and 50.9K respectively.

**Figure 1.**
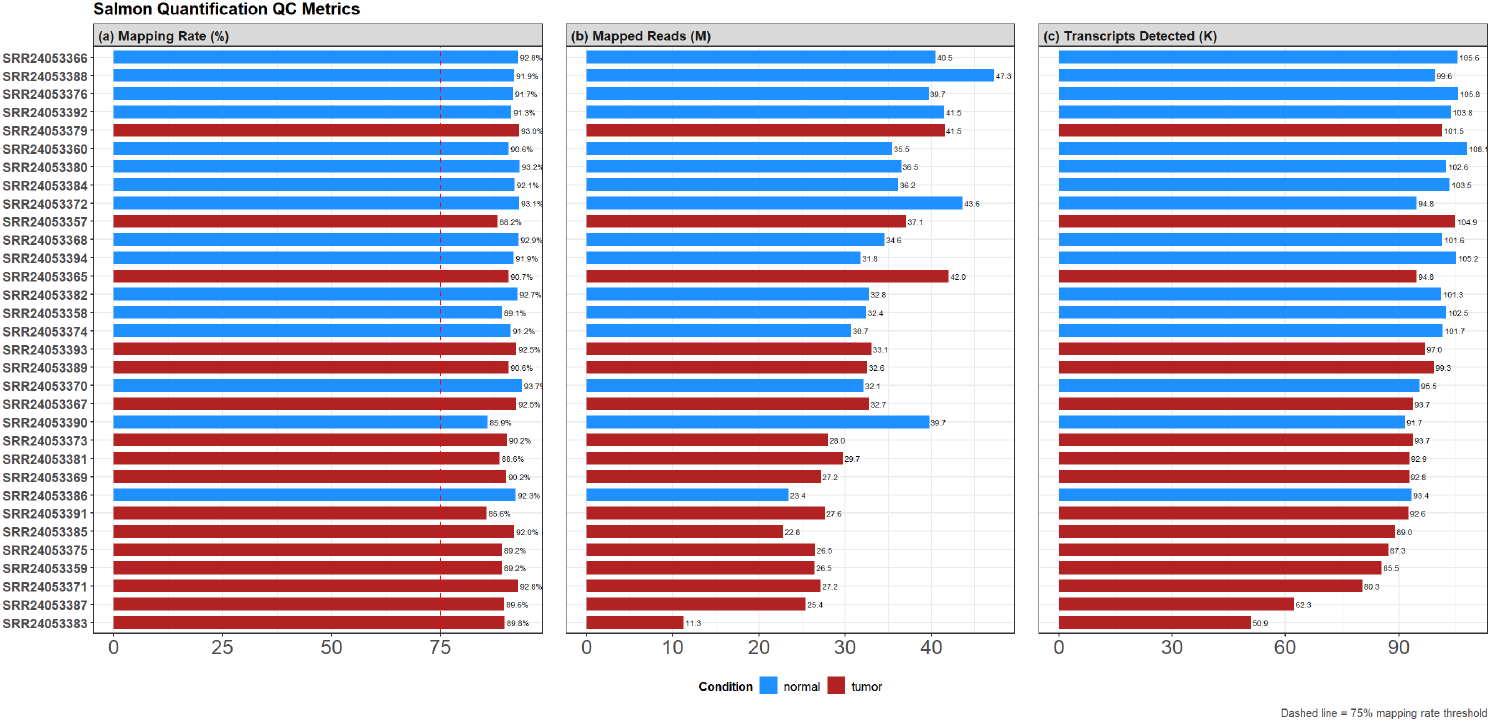
Salmon quantification quality control metrics for all 36 samples. Bar plots show (a) mapping rate (%), (b) total mapped reads (millions) and (c) number of detected transcripts (thousands) per sample. Blue and red bars represent normal and tumor samples respectively. Dashed red line indicates the 75% mapping rate threshold. Two samples (SRR24053377 and SRR24053395) falling below threshold were excluded from downstream analysis.

For SRR24053383, while the million mapped reads (11.28M) and transcript count (50.9K) were lower than other samples, its mapping rate was high (i.e. 89.8%), indicating shallow sequencing depth rather than sample contamination. The importIsoformExpression() function of *IsoformSwitchAnalyzeR()* accounts for inter-library depth variation, hence this pair was retained for downstream switch analysis. The complete QC metrics can be found in Supplementary Table S3.

### Isoform Switch Analysis Results

Following the import of Salmon quantification files, 58838 (23.45%) isoforms were removed since they were not expressed in any sample. Prefiltering was performed to remove single isoform genes and genes lacking consequence potential. This prefiltering removed 117675 (61.33%) of transcripts, leaving behind 74210 isoforms across 32 samples for downstream analysis.

PCA on log_2_-transformed TPM values from 25% most variable isoforms showed separation between normal and tumor samples (Figure 2a). Tumor samples clustered predominantly on left side of the plot while normal samples clustered on the right side. PC1 explained 37.4% of total variance and separation of tumor and normal samples along the x-axis suggested that biological condition is the primary driver of transcriptomic variability. PC2 explained 13.3% variance and captured inter-patient heterogeneity. However, samples HC004, HC005 and HC007 showed larger distances across both PCs, suggesting greater transcriptomic divergence between tumor and normal tissues in these patients. Sample HC022 showed intermediate variance across PC1 suggesting lower transcriptomic divergence between its tumor and normal samples compared to the rest of dataset.

**Figure 2.**
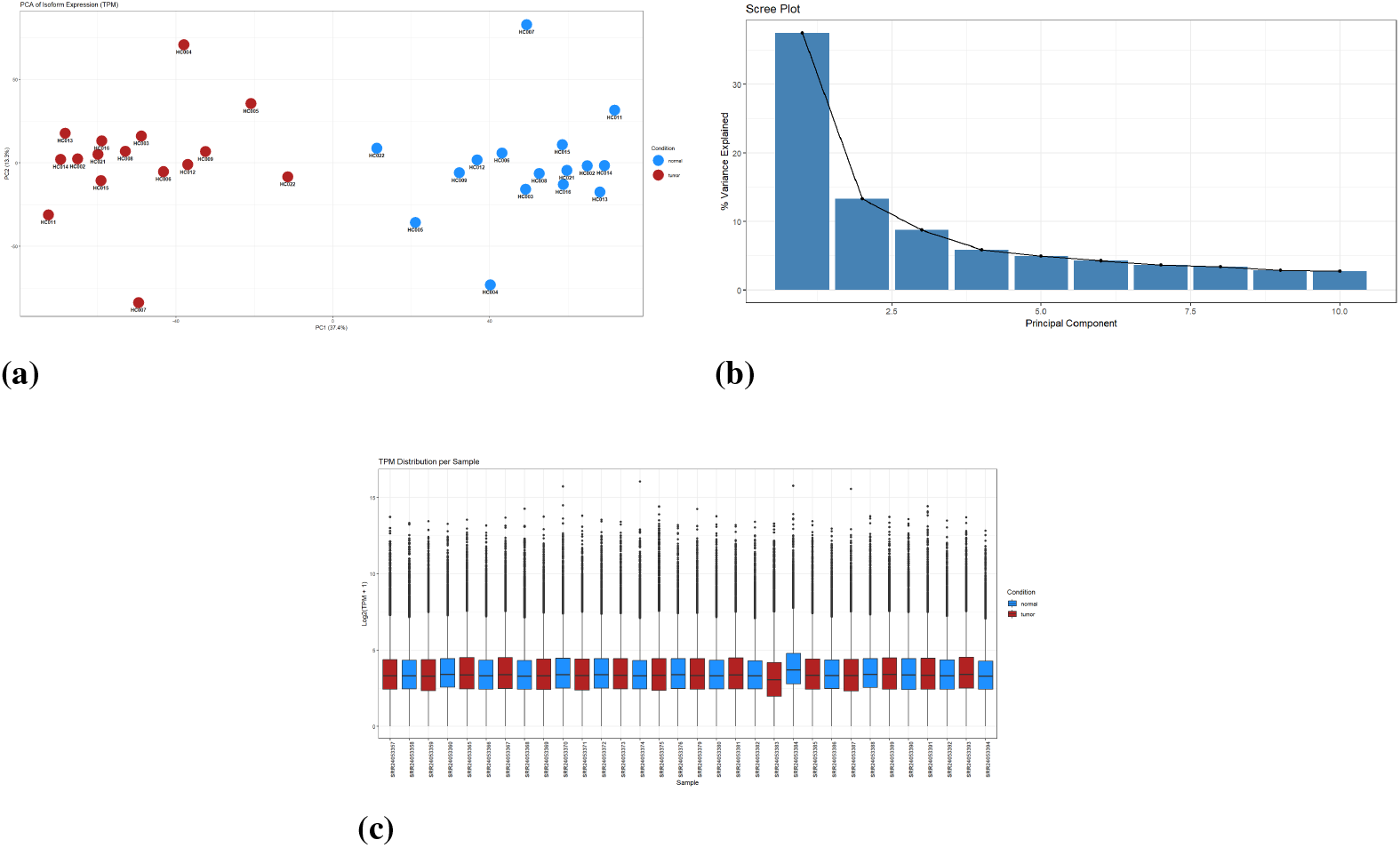
Quality control metrics for expression and normalization across samples. (**a**) PCA plot of log_2_-transformed TPM values across the top 25% most variable isoforms. (**b**) Scree plot showing percentage of variance explained by the first ten PCs. (**c**) log_2_-transformed TPM distribution (log_2_(TPM + 1)) across all 32 retained samples, confirming successful inter-library normalization.

Scree plot (Figure 2b) showed that PC1 and PC2 explained more than 50% of the cumulative variance within data, with a gradual drop after PC3. log_2_-transformed TPM values (TPM+1) were plotted for all 32 samples (Figure 2c). The boxplots showed that median and overall ranges of expression were uniform across all samples verifying the inter-library normalization of importIsoformExpression() function. For SRR24053383, the median TPM was slightly lower than other samples. This difference can be explained due to its lower sequencing depth (11.28 million mapped reads). The isoform expression profile of SRR24053383 was consistent with other retained samples for highly expressed transcripts, therefore this sample was retained for downstream analysis.

IsoformSwitchTestDEXSeq() identified 514 isoform switching events across 472 genes involving 760 distinct isoforms. ExtractSequence() function extracted 2683 nucleotide and 1600 amino acid sequences. The number of extracted amino acid sequences is lower than nucleotide number because many transcripts are non-coding or undergo NMD and do not translate to proteins. Only isoforms with annotated CDS from Gencode v44 GTF were considered for ORF analysis. These amino acid and nucleotide sequences were used for functional characterization such as prediction of coding potential, protein domain identification, IDR, sub-cellular localization, and topology information. Coding potential was assigned to 2683 (100%) transcripts via CPC2. Domain information from Pfam was added to 1399 (52.14%) transcripts. Pfam results showed that approximately half of protein isoforms map to known functional domains. The incomplete domain coverage explains the presence of non-coding transcripts lacking protein sequences for domain annotation and a subset of coding isoforms may have novel and/or truncated protein domains. IDR information from IUPred2A was added to 718 (26.76%) transcripts. IUPred2A analysis was run exclusively on amino acid sequences. IUPred2A results showed that not all proteins contain detectable disordered regions above specified thresholds. Signal Peptide information from SignalP was added to 208 (7.75%) transcripts, showing that only 7.75% of proteins contain signal peptides indicating involvement of secretory pathways. Subcellular and topology information from DeepLoc2 and DeepTMHMM was added to 1551 (57.81%) and 1600 (59.63%) transcripts respectively. DeepLoc2 predicted the sub-cellular localization of isoforms including nucleus, cytoplasm, extracellular space while DeepTMHMM predicted the number and orientation of transmembrane helices. Both tools predicted sub-cellular localization and transmembrane topology for more than half the analyzed transcripts. Alternative splicing analysis revealed that 419 isoforms exhibited intron retention events, with majority of isoforms retaining at least one intron (n = 322).

Differential isoform usage identified 514 statistically significant isoform switches across 472 genes as significant (*q*-value < 0.05 and |*dIF*| > 0.1). After filtering the list for functional consequences, 371 isoform switches across 335 genes involving 551 isoforms remained. Switch consequence summary plot (Figure 3) shows functional outcome of isoform switches in HCC. The plot indicates that isoforms in HCC lose protein domains more frequently than gain protein domains (domain gain = 132 vs domain loss = 150) and showed reduced topology complexity (topology complexity gain = 99 vs topology complexity loss = 139). Moreover, isoforms in HCC resulted in production of shorter ORFs (shorter ORF = 138 vs longer ORF = 125) and 106 isoforms retained their coding potential compared to 89 non-coding ORFs. Isoforms expressed in HCC frequently exhibited signal peptide loss (n=37) and are expressed in different sub-cellular localization than their normal counterparts (n=28). Complete consequence switch statistics can be found in Supplementary Table S4. Collectively, these findings indicate that isoforms overexpressed in HCC encode for shorter ORFs with smaller protein domains, have simpler membrane topology, reduced involvement in secretory pathways and are predicted to acquire different sub-cellular localizations than their normal counterparts.

**Figure 3.**
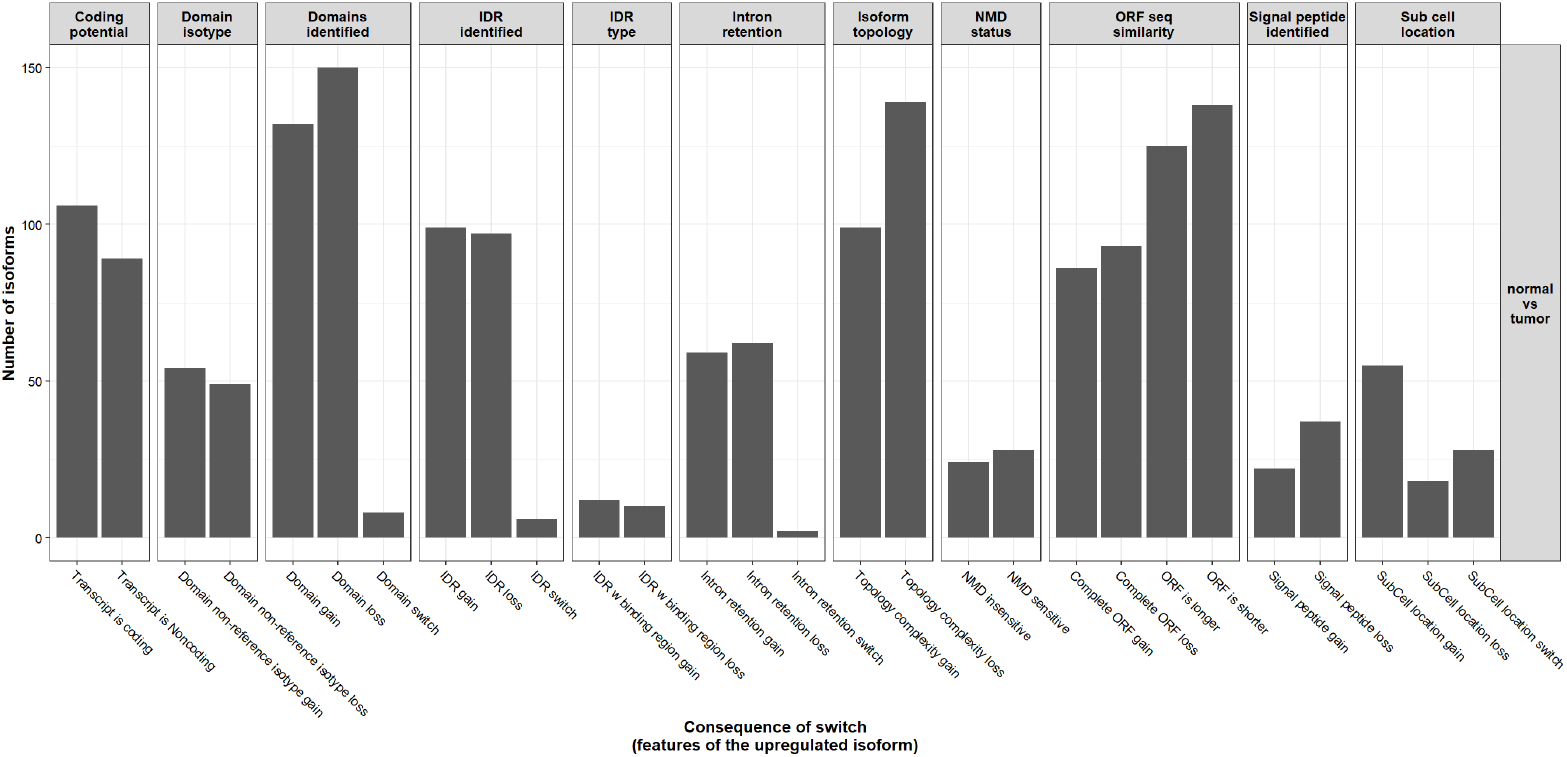
Summary of functional consequences of isoform switches in HCC. Bar plots show the number of upregulated isoforms exhibiting each consequence type across eleven functional categories including coding potential, protein domain content, IDR, IR, topology, NMD sensitivity, ORF similarity, signal peptide, and subcellular localization.

Top genes filtered after *q*-value and |dIF| cutoff included *LAMA2* (rank = 1, |dIF| = 0.197, *q*-value = ×10^−31^), *LSP1* (rank = 2, |dIF| = 0.386, *q*-value = 5.27 × 10^−20^), *MAD2L2* (rank = 3, |dIF| = −0.331, *q*-value = 7.12 × 10^−19^), *FBLN2* (rank = 8, |dIF| = 0.361, *q*-value = 3.94 × 10^−11^) and *CXCL12* (rank = 10, |dIF| = −0.334, *q*-value = 4.31 × 10^−11^). Complete information about the isoform, domain architecture, single peptides, NMD status and sub-cellular localization of these genes can be found in Table 1. These genes are known to be involved in thyroid carcinoma. The genes ranked between 4 to 7 i.e., *MBD6*, *MPZ*, *PEX19* and *VAPB* are not known to be involved in any thyroid malignancy and hence were not explored for interpretation. A complete list of top 20 significant isoforms along with their metrics can be found in Supplementary Table S5. Switch plots for all these genes were used to visualize the structure of individual isoforms, compare overall gene and isoform expression and relative isoform usage in controls and HCC samples.

**Table 1.** Top 5 most significant isoform switches in HCC filtered by *q*-value < 0.05 and |dIF| > 0.1 with confirmed functional consequences.

| Rank | Gene | Isoform | dIF | q-value | Domains Identified | IDR | Signal Peptide | NMD Status | Sub-cellular Localization |
| --- | --- | --- | --- | --- | --- | --- | --- | --- | --- |
| 1 | *LAMA2* | ENST00000617695.5 | 0.197025 | 1.08E-31 | Yes | No | Yes | False | Extracellular |
| 2 | *LSP1* | ENST00000311604.8 | 0.386013 | 5.27E-20 | Yes | Yes | No | False | Cytoplasm |
| 3 | *MAD2L2* | ENST00000376692.9 | -0.33106 | 7.12E-19 | Yes | No | No | False | Cytoplasm |
| 8 | *FBLN2* | ENST00000295760.11 | 0.361494 | 3.94E-11 | Yes | Yes | Yes | False | Extracellular |
| 9 | *FBLN2* | ENST00000404922.8 | -0.36071 | 3.94E-11 | Yes | Yes | Yes | False | Extracellular |
| 10 | *CXCL12* | ENST00000343575.11 | -0.33413 | 4.31E-11 | Yes | No | Yes | False | Extracellular |

Switch Plot for *LAMA2* (Figure 4a) showed that the overall gene expression of *LAMA2* is reduced to half in tumor tissues, while the absolute expression of one if its isoforms ENST00000617695.5 is increased in tumor tissues. Isoform usage plot showed that ENST00000617695.5 is used significantly more in HCC than in normal tissues. On the other hand, usage of ENST00000421865.3 and ENST00000688799.1 is significantly decreased in tumor tissues relative to normal tissue. ENST00000617695.5 is one of the coding isoforms of *LAMA2* and the switch plot show that the isoform lacks a C-terminus protein region present in the canonical isoform ENST00000421865.3. This truncation may compromise the laminin polymerization within extracellular matrix (ECM). ENST00000688799.1 is also a coding isoform and has relatively simpler structure consisting of shorter sequences and fewer domains than other isoforms of *LAMA2*.

**Figure 4.**
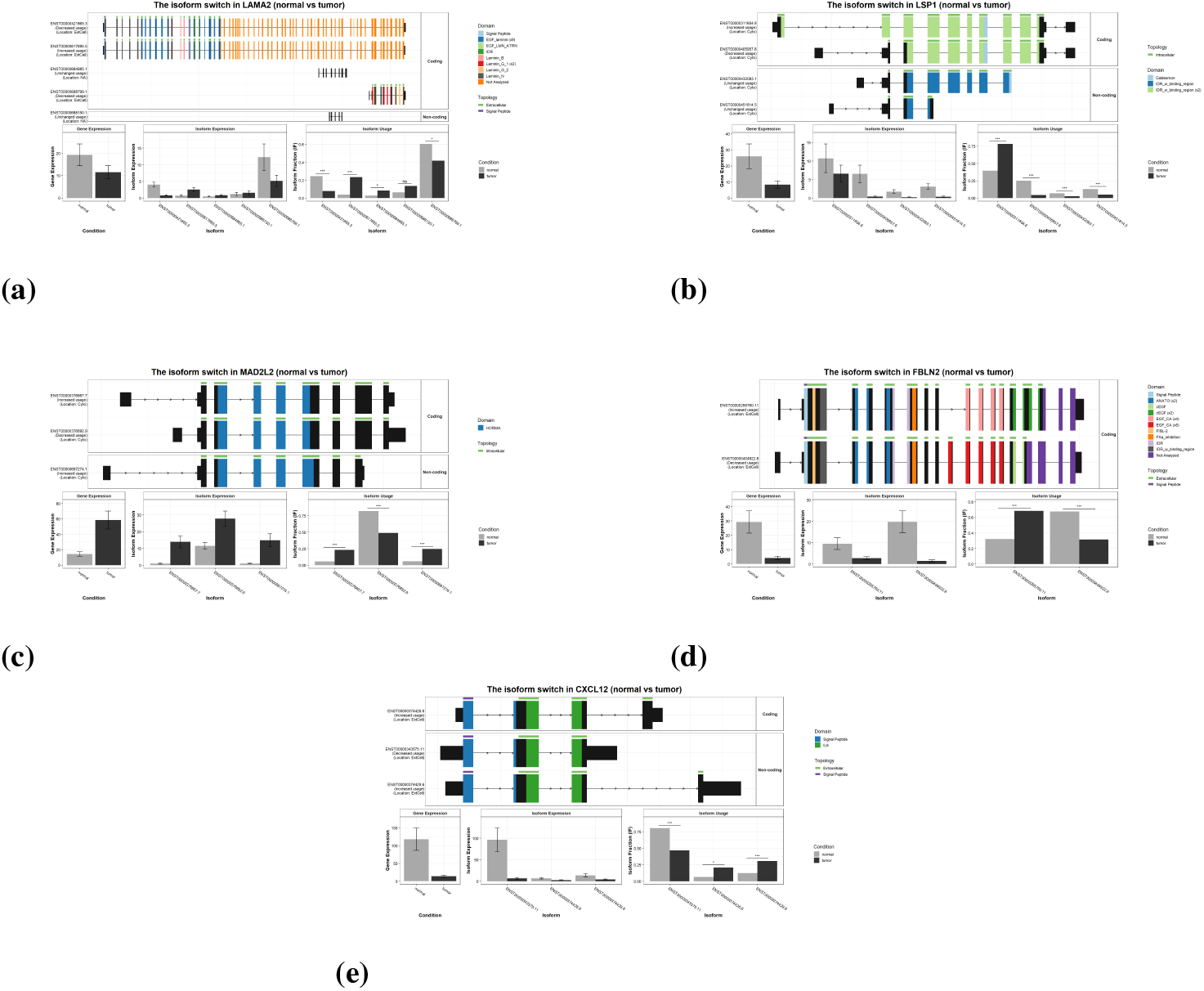
Isoform switch plots for key genes in HCC. For each panel, the top section shows transcript structures with annotated protein domains and topology; the bottom section shows gene expression, absolute isoform expression, and relative isoform usage in normal and tumor conditions. (**a**) Plot for *LAMA2*, demonstrating ENST00000617695.5 lacks a C-terminus protein region. (**b**) Plot for *LSP1*, demonstrating predominant shift toward the Caldesmon-containing isoform in tumor. (**c**) Plot for *MAD2L2* with HORMA domain annotation. (**d**) Plot for *FBLN2*, demonstrating shift toward four tandem repeat CA-binding EGF containing isoform in tumor. (**e**) Plot for *CXCL12*, demonstrating increased usage of truncated coding and non-coding isoforms in tumor.

Switch plot for *LSP1* (Figure 4b) indicates that absolute gene and isoform expression is reduced in HCC tissues compared to normal cells. However, the relative expression of isoform ENST00000311604.8 is significantly increased in tumor cells. ENST00000311604.8 is a coding isoform composed of caldesmon domain and two IDR binding regions. ENST00000311604.8 has an extended first exon and additional two IDR binding regions at 5’ end. To summarize, the overall *LSP1* expression is reduced in HCC but the relative usage of one of its coding isoforms is increased. Caldesmon and IDR binding regions within coding isoform suggest an altered cytoskeletal regulatory capacity in HCC.

Switch plot for *MAD2L2* (Figure 4c) shows significant overexpression of this gene in HCC tissues. The absolute expression of all its isoforms (ENST00000376667.7, ENST00000376692.9 & ENST00000697274.1) is significantly increased in tumor cells. However, the relative expression of ENST00000376667.7 and ENST00000697274.1 is increased while ENST00000376692.9 showed markedly reduced usage in HCC. ENST00000376667.7 is a coding isoform composed of intracellular HORMA domains and is seen to have an extended 5’ region as compared to the other coding isoform ENST00000376692.9. The other coding isoform, ENST00000376692.9, which has significantly decreased usage in tumor cells (IF drops to approximately 0.48) have a truncated 5’ and 3’ exon. The non-coding isoform of *MAD2L2*, ENST00000697274.1, shows shorter 3’ UTR/terminal exons and a shorter 5’ exon. The expression of *MAD2L2* in HCC changes from single coding isoform to a mixed population of both coding and non-coding isoforms. Notably, the HORMA domain, which is required for the DNA damage response, is successfully retained in upregulated coding isoform ENST00000376667.7, suggesting that despite the isoform usage shift, domain-level functional capacity of *MAD2L2* may be preserved in HCC.

The absolute expression of *FBLN2* (Figure 4d) gene and its isoforms is found to be down regulated in HCC compared to normal tissue. However, the usage of one of its isoforms ENST00000295760.11 is significantly increased in tumor (dIF=0.36) and the other isoform ENST00000404922.8 is significantly used less in HCC than normal by the same amount (dIF=-0.36). Both isoforms of *FBLN2* are coding, but the switch plot shows several structural differences between them. The isoform with increased usage (ENST00000295760.11) shows four tandem repeats of extracellular calcium binding (CA) epidermal growth factor (EGF) domains (EGF CAx4) compared to its counterpart (ENST00000404922.8) which shows presence of five tandem repeats of extracellular CA EGF-binding domains (EGF CAx5). ENST00000295760.11 shows an additional two tandem repeat cEGF domain while this tandem repeat is absent in ENST00000404922.8. Additionally, ENST00000295760.11 shows extended 3’ terminal exon. Collectively, gene-level downregulation coupled with switch towards an isoform of *FBLN2* with reduced EGF CA repeat in HCC suggests its involvement in HCC ECM disruption.

The overall gene expression of *CXCL12* (Figure 4e) is significantly reduced in tumor cells than normal. Similarly, the absolute expression of all its isoforms (ENST00000343575.11, ENST00000374426.6 & ENST00000374429.6) is also significantly reduced in tumor tissues. Contrastingly, the isoform usage plot shows that relative expression of two of its isoforms (ENST00000374426.6 & ENST00000374429.6) is increased in tumor tissues. ENST00000374426.6 is a coding isoform composed of signal peptide and extracellular interleukin-8 (IL8) domains, with truncated 5’ and 3’ ends. ENST00000374429.6 is a non-coding isoform and is composed of relatively elongated 3’ UTR/terminal exon. Collectively, *CXCL12* in HCC showed gene-level downregulation as well as a proportional shift towards production of truncated, non-coding isoform.

A summary graph of overall alternative splicing events for 760 isoforms (Figure 5) showed that ATSS was the dominant event in both isoforms used more (n=352 isoforms, 338 genes) and less in HCC (n=346 isoforms, 338 genes). ATTS showed a directional asymmetry, with isoforms gaining this event in tumor (n=312 isoforms, 296 genes), thereby explaining shift towards alternative termination site usage in HCC. ES occurred at comparable frequencies in isoforms used more (n=223) and less (n=245), with a slight bias towards isoforms used less. MES showed similar bidirectional occurrence (used more: n=102; used less: n=115). IR contributed modestly in both directions (used more: n=57; used less: n=59). MEE showed negligible contribution (used more: n=3; used less: n=1). Complete alternative splicing events statistics across isoforms can be found in Supplementary Table S6

**Figure 5.**
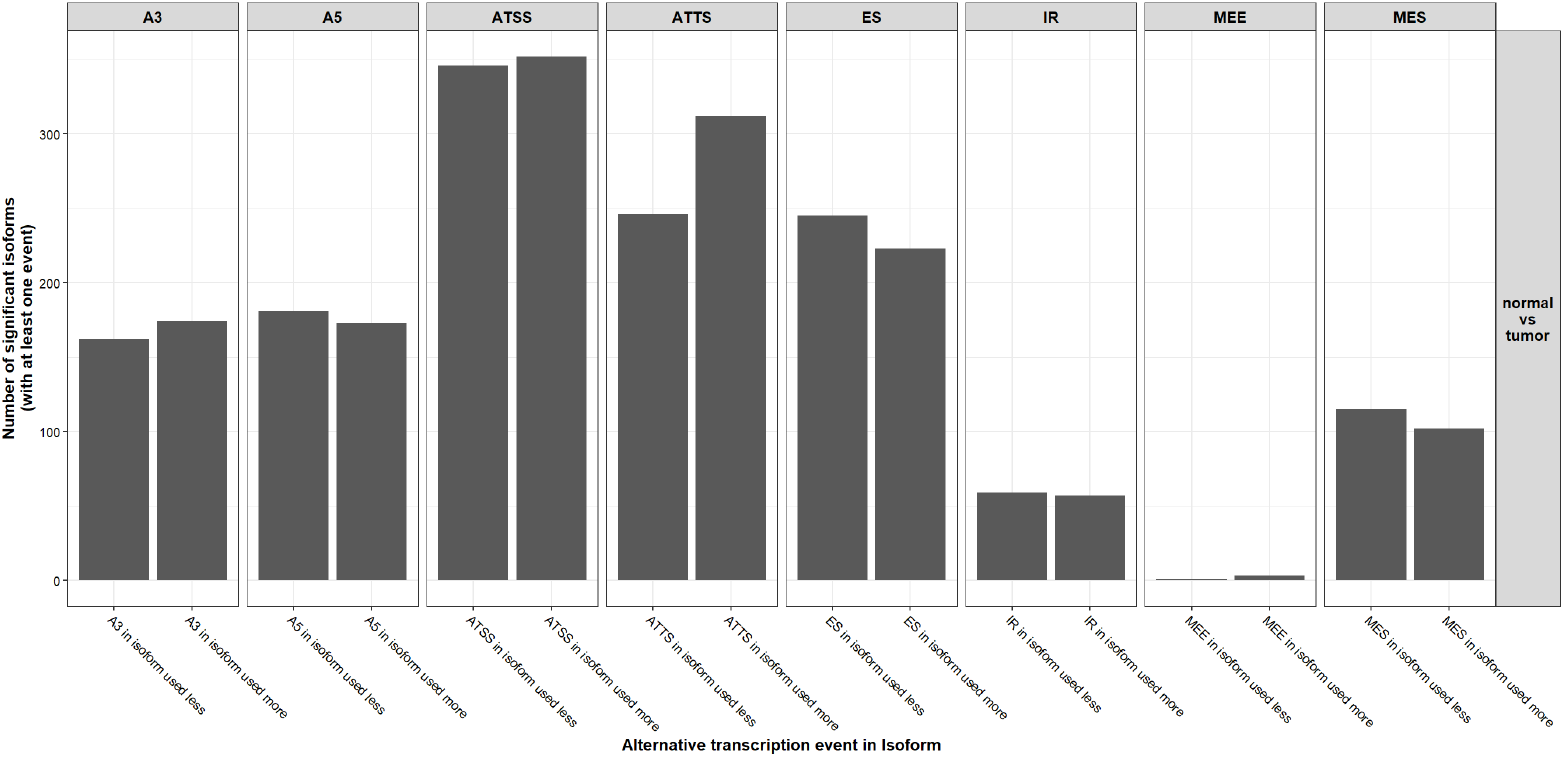
Summary of alternative splicing events across 760 significant switching isoforms. Bar plots show the number of isoforms exhibiting each event type (A3, A5, ATSS, ATTS, ES, IR, MEE, MES), stratified by direction of isoform usage change (used more vs. used less) in HCC relative to normal tissue.

Among the switches with confirmed functional consequences, AS, ATSS and ATTS were evaluated. All three events had almost the same occurrence (AS: 266, ATTS: 254 & ATSS: 244). This shows that tumor cells utilize multiple mechanisms to produce isoforms. Splice enrichment graph (Figure 6) was used to visualize the statistical significance of each splicing event type. The graph showed that ATTS gain was statistically significantly enriched than ATTS loss (*q*-value < 0.05). This shows that isoform switches in HCC preferentially involves a shift towards transcripts with altered alternative 3’ ends, suggesting dysregulation of transcript termination as the mechanism behind isoform diversity in HCC. Complete results of splicing enrichment analysis can be found in Supplementary Table S7.

**Figure 6.**
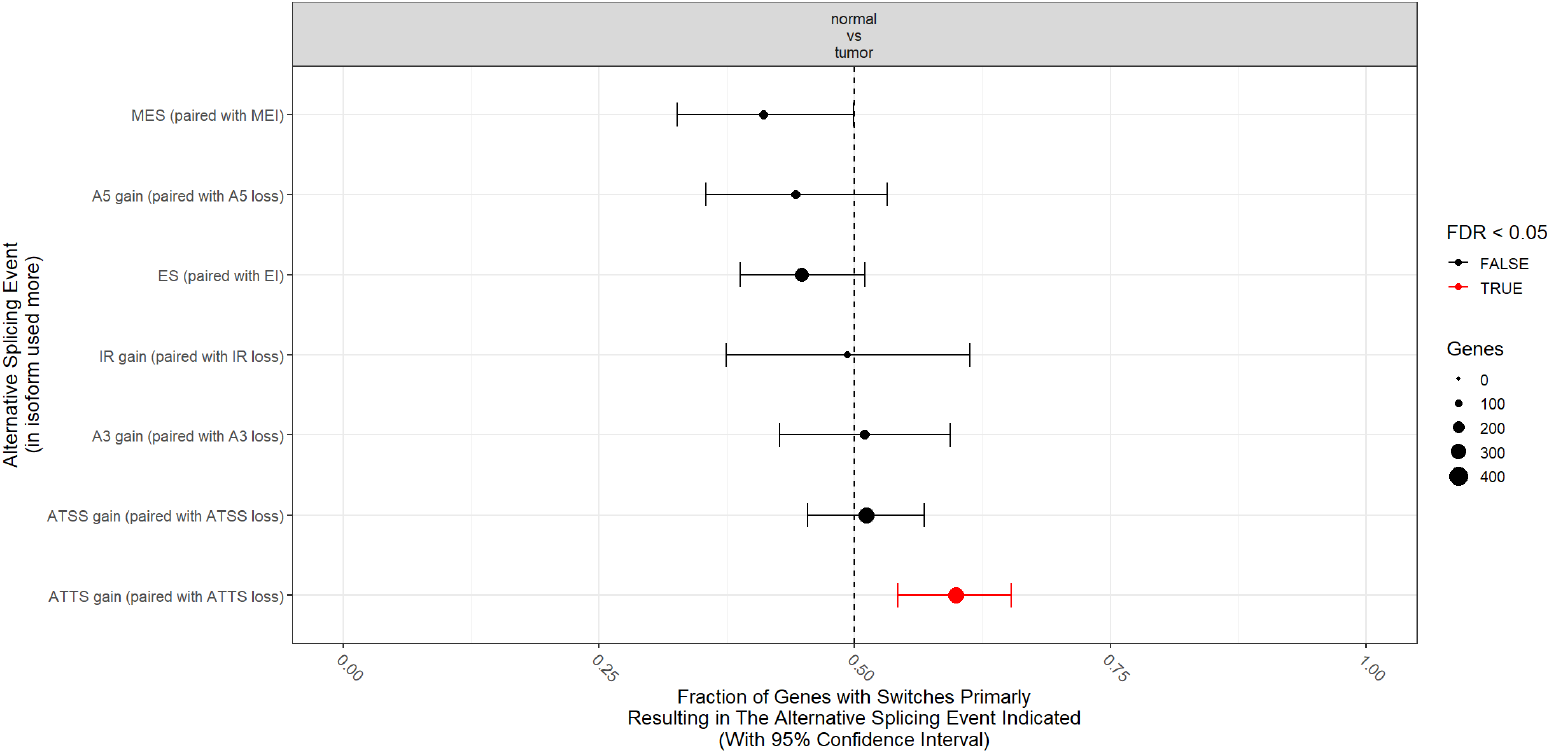
Splicing enrichment analysis showing directional bias of alternative splicing events in HCC. Each point represents the fraction of genes where switches resulted in gain of the indicated splicing event, with 95% confidence intervals. Red point indicates statistically significant enrichment (*q*-value < 0.05). Dot size reflects the number of contributing genes.

### Pathway Enrichment Results

The 335 genes showing isoform switching with functional consequences in HCC were used to perform functional enrichment analysis. Our target genes including *LAMA2*, *LSP1*, *MAD2L2*, *FBLN2* and *CXCL12* showed involvement in molecular functions including protein binding (GO:0005515, *q*-value = 2.54 × 10^−7^) and extracellular matrix structural constituents (GO:0005201) (Figure 7). Moreover, enrichment analysis showed that *LAMA2*, *MAD2L2*, *FBLN2* and *CXCL12* are involved in biological process such as lymphocyte activation (*q*-value = 0.0100), regulation of primary metabolic processes (*q*-value = 0.020), in regulation of cell development (*q*-value = 0.022), cell motility (*q*-value = 0.023), and regulation of transcription (*q*-value = 0.026). Complete results from g:Profiler can be found in Supplementary Table S8.

**Figure 7.**
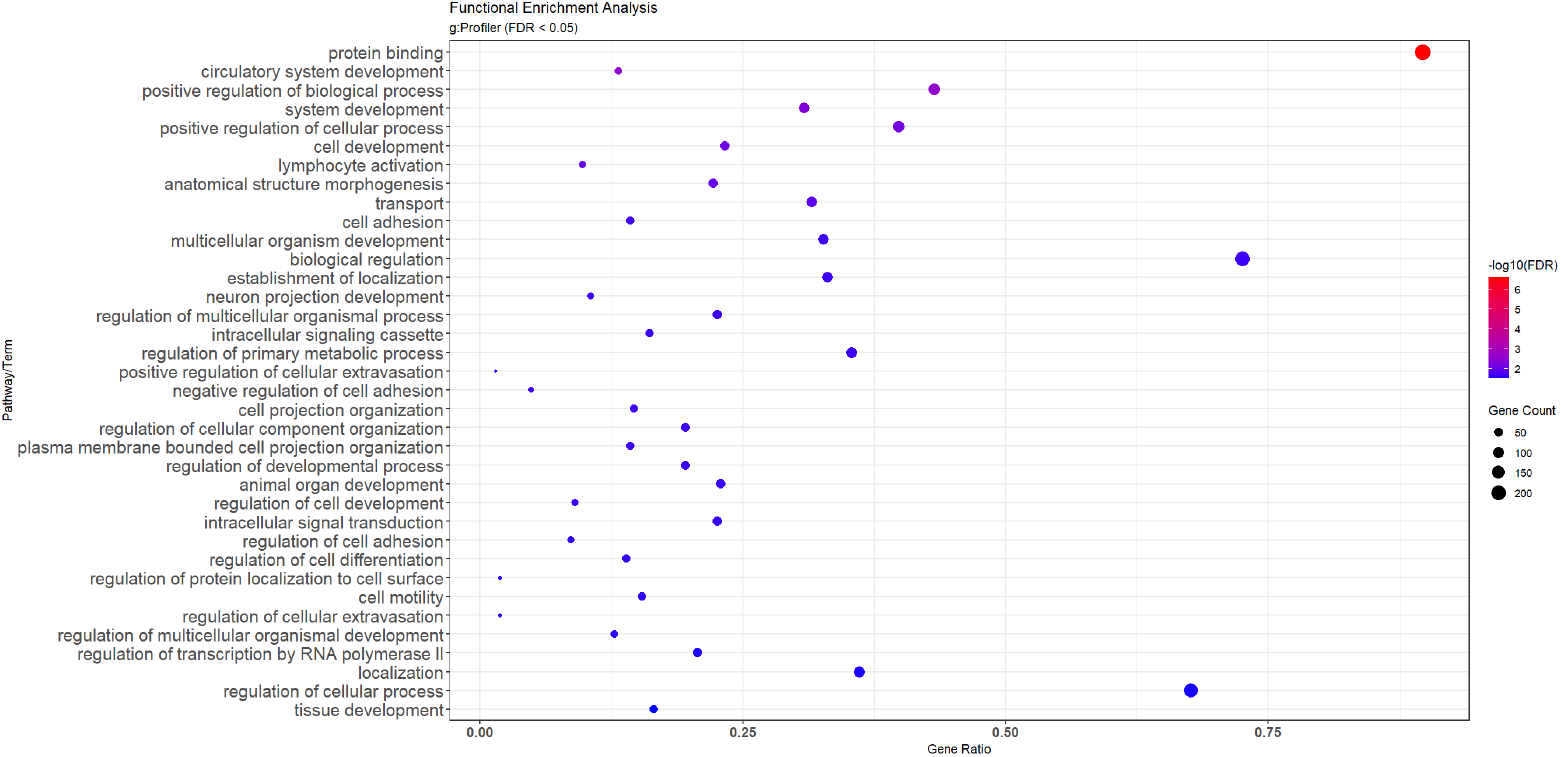
Dot plot showing enriched molecular functions and biological processes in HCC. Each dot represents the fraction of genes involved in a process, with *q*-value < 0.05. Dot size reflects the number of contributing genes.

### GTEx and TCGA SpliceSeq Validation Results

GTEx portal showed that all top 10 genes undergoing alternative splicing in HCC are expressed in normal thyroid tissue (Table 2). *LAMA2* isoform with increased usage in HCC (ENST00000617695.5) had relatively low expression in normal thyroid (0.170 TPM) suggesting a tumor-associated shift in HCC. For *LSP1*, isoform ENST00000311604.8 showed highest expression in normal thyroid tissue (10.9 TPM) followed by ENST00000405957.6 (6.86 TPM). However, differential isoform usage calculated by *IsoformSwitchAnalyzeR()* showed that ENST00000311604.8 is used more in HCC than normal tissue. As ENST00000311604.8 is expressed in both normal thyroid and HCC, its increased usage in tumor suggests pathogenic relevance in HCC. *IsoformSwitchAnalyzeR()* results showed that expression of all *MAD2L2* isoforms is relatively high in HCC than normal tissue. GTEx results also show that the isoforms with increased usage in HCC (ENST00000376667.7 & ENST00000697274.1) do not show any expression in normal thyroid tissue. The absence of ENST00000376667.7 and ENST00000697274.1 in normal tissue explains that these isoforms are tumor specific and their increased usage represents aberrant splicing regulation in HCC. For *FBLN2* isoform ENST00000295760.11 is expressed low in HCC, but its differential usage is increased in tumor tissues suggesting tumor-associated shift in isoform preference. *CXCL12* isoforms used more in HCC are also expressed in normal thyroid, suggesting a tumor-associated shift in isoform preference.

**Table 2.** Summary of GTEx and TCGA SpliceSeq validation results.

| Gene | GTEx Median TPM in normal thyroid tissue (Sample Size = 684) | HCC Isoform | Isoform TPM in normal thyroid | TCGA SpliceSeq even type | PSI Value (SpliceSeq) | Summary of *IsoformSwitchAnalyzeR()* Results |
| --- | --- | --- | --- | --- | --- | --- |
| LAMA2 | 20.69 | ENST00000688799.1 | 33.1 | ES | 49.1% for exon44<br>42.2% for exon53 | Low expression in normal & increased usage in HCC |
| LSP1 | 11.66 | ENST00000311604.8 | 10.9 | AP & ES | 94.5% for exon1 (AP)<br>95.5% for exon7 (ES) | High expression in normal but increased usage in HCC |
| MAD2L2 | 14.79 | ENST00000376667.7 | 0 | ES | 82.6% for exon3 | Absent in normal thyroid, tumor-specific expression |
| FBLN2 | 134.1 | ENST00000295760.11 | 35.3 | AP & ES | 99% for exon2 (AP)<br>96.3% for exon12 (ES) | Low expression but increased usage in HCC |
| CXCL12 | 80.76 | ENST00000374429.6 | 18.5 | ES & RI | 96.3% for exon2.2 (ES)<br>99.8% for exon3.2 (RI) | Low expression but increased usage in HCC |

Splicing mechanisms for top 10 genes were validated through TCGA SpliceSeq by applying thyroid cancer filter. SpliceSeq showed that *LAMA2*, *MAD2L2*, *LSP1*, *FBLN2* and *CXCL12* undergo ES in thyroid cancer explaining the truncated domain structure observed in isoform ENST00000617695.5, ENST00000376667.7, ENST00000697274.1 and the change of EGF-binding domains tandem repeats in isoform ENST00000295760.11. Additionally, *LSP1* show alternative promoter (ATSS) explaining the extended 5’ region observed in ENST00000311604.8. *CXCL12* show IR and alternative termination (ATTS) which would explain the elongated 3’ UTR of CXCL12’s ENST00000374429.6 isoform.

## Discussion

This is the first study which characterizes functional isoform switches and their consequences in context of HCC. Through this analysis, we identified 514 switching events across 472 genes, resulting in 760 distinct isoforms. Among these, 551 isoforms showed functional consequences including loss of protein domains, reduced topology complexity, shorter ORFs, reduced involvement in secretory pathways and novel sub-cellular localizations. Most significant isoform switches (*q*-value < 0.05 and |dIF| > 0.1) were observed in *LAMA2*, *LSP1*, *MAD2L2*, *FBLN2* and *CXCL12* genes, suggesting involvement of ECM dysregulation, DNA damage response, immune signaling and cytoskeleton regulation. ATTS gain was statistically significant, suggesting a shift towards altered 3’ end in transcripts expressed in HCC.

Previous studies have explored the landscape of isoform switching in different solid tumors such as brain, breast, kidney, liver, thyroid, ovary, prostate and skin. These studies have extensively analyzed the functional impact of isoform switches on protein domains. One such study analyzed isoform switches in 12 different cancer types using data from TCGA and reported that annotated proteins in tumor isoforms are shorter than proteins encoded by isoforms in normal tissues. The authors also confirmed that tumor isoforms are associated with loss of protein coding regions. Moreover, normal isoforms tend to be longer than tumor isoforms (Climente-González et al. 2017). Analysis of functional consequence of isoform switches in 12 different cancer types reported that upregulated isoforms in cancer lead to significant loss of protein domains. ORF analysis showed that cancer isoforms have significant overrepresentation of ORF loss, coupled with gain of signal peptide (Vitting-Seerup and Sandelin 2017). While our study reported loss of protein domains, and shorter ORFs, we observed loss of signal peptides in HCC isoforms. This suggests tumor specific differences exist with regard to secretory pathway regulation. Pathogenic impact of isoform switches in 27 different cancers reported that isoforms upregulated in cancer have disruptions in protein translation and transcription. Isoform switching results in loss of kinase domains in proteins (Kahraman et al. 2020). Pan-cancer isoform analysis showed that cancer-associated isoforms show distinct expression patterns from their normal tissue counterparts, consistent with domain loss and structural simplification observed in HCC isoforms in current analysis (Surana et al. 2026).

AP or ATTS is a known dysregulated mechanism in cancer. 3’ UTR shortening through AP is known to promote tumor growth (Park et al. 2018; Gabel et al. 2024). However, conflicting studies have been observed in this regard. Some studies report shortening of 3’ UTR in cancer isoforms while others show lengthening of 3’ UTR in cancer. Colorectal adenocarcinoma patients have reported long 3’ UTR coupled with loss-of-function adenomatous polyposis coli (*APC*) mutations (Gabel et al. 2024). Contrastingly, breast cancer patients show shortening of 3’ UTR which can contribute to cancer growth in a subtype-specific manner (Fan et al. 2020). Shortening of 3’ UTR destabilizes mRNA of oncogenes and in turn enhances the stability of tumor suppressor genes’ mRNA in clear cell renal cell carcinoma (ccRCC). In ccRCC, lengthening of 3’ UTR is associated with tumor aggressiveness and heterogeneity (Zhang et al. 2023). Proliferating cells, including cancer cells, are known to give rise to isoforms with shorter 3’ UTR, owing to cleavage and polyadenylation at proximal coding region sites. Subsequently, isoforms with 3’ UTR are less susceptible to post-transcriptional regulation due to loss of microRNA (miRNA) binding sites. Hence, change in 3’ UTR length contributes to progression and prognosis of cancer (Schmidt et al. 2018). A study analyzing isoform switch consequences in 12 different cancer types showed that AS and ATSS were the major regulatory mechanisms controlling the production of isoforms. On the other hand, ATTS only contributed to 17.3% switches (Vitting-Seerup and Sandelin 2017). As our analysis reports ATTS as the major mechanism of alternative splicing in HCC tumor isoforms, we propose that ATTS gain is tumor specific splicing mechanism and potentially contributes to isoform diversity through altered transcription termination.

Genes showing isoform switching in HCC have increased role in cancer development and progression. *LAMA2* gene is present on chromosome 6q22-q33 and encodes Laminin subunit *α*2. *LAMA2* is involved in ECM adhesion and have increased roles in the progression of breast cancer, ovarian cancer, pituitary adenoma and squamous cell carcinoma. Additionally, *LAMA2* have shown regulatory roles in molecular mechanisms and signaling pathways via DNA methylation (Hu and Li 2022). Methylation of *LAMA2* promoter is found to be increased in pituitary adenomas, which in turn leads to the decreased expression of *LAMA2* gene. Hence, methylation and expression pattern of *LAMA2* is proposed to act as potential biomarkers in pituitary adenoma patients (Wang et al. 2019). In HCC, an increased usage towards C-terminus truncated domain isoform suggests impaired ECM network assembly and tumor invasion, consistent with *LAMA2*’s role in cancer progression. Whether promoter methylation contribute to *LAMA2* under expression in HCC remains to be investigated.

Leukocyte-specific protein 1 (*LSP1*) is an F-actin binding protein, contains Caldesmon domains, and is expressed in macrophages, neutrophils and endothelial cells. Mutations of *LSP1* have been reported in various tumors. *LSP1* copy number variations have been reported in hepatocellular carcinoma. *LSP1* is known to function as negative regulator of proliferation and migration in mouse liver cells (Koral et al. 2014). *LSP1* rs3817198 (T>C) mutation is known to cause breast cancer malignancy among Caucasians and Asians (Chen et al. 2022). *LSP1* act as a novel prognostic biomarker of acute myeloid leukemia (AML) and increases the apoptosis of CD8+ T cells in AML (Xu et al. 2025b). *LSP1* is reported to be significantly increased in cervical cancer tissues and correlates with better patient outcomes (Xu et al. 2025a). Contrastingly, *LSP1* was found to be under expressed in HCC with one of its isoforms being significantly used more than others. However, the isoform used more in HCC showed retention of Caldesmon domain suggesting conserved domain structure of *LSP1*.

Mitotic arrest deficient 2 like 2 (*MAD2L2*) is a chromatin binding protein and is a part of DNA polymerase *ζ*. *MAD2L2* is part of shieldin complex and plays a role in DNA double strand break repair (Krijger et al. 2021). *MAD2L2* is known to be involved in cell cycle regulation, DNA mismatch repair and cancer progression (Krijger et al. 2021). Genetic alterations including amplification and deletions within *MAD2L2* and overexpression of *MAD2L2* is associated with increased cell proliferation, migration and reduced survival rates in ovarian cancer patients, particularly with those in grade IV tumors (Xu et al. 2024). The overall gene expression of *MAD2L2* is found to be increased in HCC, which is consistent with its reported role in ovarian cancer, suggesting a potentially similar cancer progression mechanism. *MAD2L2* contains conserved HORMA domain essential for its dimerization and protein-protein interaction (Barda et al. 2024). The isoform showing increased usage in HCC have retained HORMA domain suggesting that despite isoform switching, the structural functionality of *MAD2L2* in HCC is preserved.

Fibulin-2 (*FBLN2*) is a large calcium-binding ECM protein which is expressed in normal epithelia. *FBLN2* contains a tandem repeat of EGF domains coupled with C-terminal and N-terminal fibulin-type modules. *FBLN2* shows both oncogene and tumor suppressor roles dependent on cancer type. In intestinal and respiratory tumor cells, its downregulation has been associated with cancer development (Zhang et al. 2020). *FBLN2* is required to maintain basal membrane (BM) integrity in mammary epithelia and loss of *FBLN2* is associated with breast cancer invasion (WalyEldeen et al. 2024). Splicing dysregulation in *FBLN2* is associated with colorectal cancer invasion. Colorectal cancer patients show exclusion of exon9 of *FBLN2*, which results in loss of single N-glycosylation site leading to misfolding of *FBLN2* protein. This misfolded *FBLN2* protein has low stability and secretion efficiency. Moreover, in colorectal cancer, down regulation of *FBLN2* is associated with changes in tumor microenvironment leading to invasion and metastasis (Funayama et al. 2025). The reduction from five to four tandem repeats of extracellular calcium-binding EGF domains in HCC isoforms may compromise the regulatory role of *FBLN2* in BM ans suggests tumor-specific role of *FBLN2* in HCC.

Chemokine (CXC motif) ligand 12 (*CXCL12*) belongs to CXC subfamily of chemokines which binds to its receptor CXC chemokine receptor 4 (*CXCR4*) or *CXCR7* and mediates cell proliferation, cell adhesion, survival and cell migration. *CXCL12* is known to be overexpressed in various cancers including pancreatic cancer, breast cancer, PTC, glioblastoma and prostate cancer. In PTC, elevated levels of *CXCL12* correlate with lymph node metastasis (Chen et al. 2019). mRNA expression of *CXCL12* in PTC have been significantly correlated with large tumor size, tumor stages III and IV and metastasis. Moreover, PTC patients with extra-thyroid extension show an increased level of *CXCL12* (Attia et al. 2024). Contrastingly, another study showed that aberrant methylation of *CXCL12* promoter can lead to PTC development and progression, through under expression of *CXCL12* (Zhang et al. 2017). As our analysis reported under expression of *CXCL12* in HCC, whether promoter methylation contributes to *CXCL12* down regulation needs epigenomic investigation.

Pathway enrichment analysis showed the involvement of genes (*LAMA2*, *LSP1*, *MAD2L2*, *FBLN2*, & *CXCL12*) in protein binding, extracellular matrix structural components and regulation of primary metabolic processes. This suggests that isoform switching in these key genes may contribute to dysregulation of biological processes relevant to HCC pathogenesis. Validation through GTEx and TCGA SpliceSeq show that genes showing alternative splicing and isoform switching in HCC are also expressed in normal thyroid tissue. *LAMA2* and *MAD2L2* isoforms (ENST00000617695.5 for *LAMA2*, ENST00000376667.7 & ENST00000697274.1 for *MAD2L2*) which show increased usage in HCC are under expressed in normal thyroid, while *FBLN2*’s isoform ENST00000295760.11 is expressed low in HCC but its usage is increased in tumor. These results suggest that these isoforms are tumor-related and their increased usage in HCC represents dysregulated splicing. SpliceSeq results show that *LAMA2*, *MAD2L2*, *LSP1* and *CXCL12* undergo concordant splicing mechanisms across thyroid cancer and HCC, suggesting that splicing dysregulation is shared across thyroid malignancies. Notably, *LSP1*, *FBLN2* and *CXCL12* were reported to undergo alternative promoter (PSI: 94.5% for *LSP1*) and alternative termination (PSI: 99.0% for *FBLN2* & 53.9% for *CXCL12*) respectively confirming ATTS and ATSS as significant splicing mechanisms in HCC (Liu et al. 2024). Enrichment analysis confirmed that top genes undergoing isoform switches in HCC are involved in ECM structural composition, protein binding and metabolic regulation. These findings are consistent with the hypothesis that mitochondrial dysregulation and chromosomal instability in HCC further dysregulate downstream transcriptomic processes through isoform switching.

Alternative splicing signatures have been previously associated with overall survival (OS) of thyroid cancer patients. A study analyzing alternative splicing events in thyroid carcinoma using TCGA SpliceSeq identified ES, AP and AT as events associated with poor prognosis (Wu et al. 2021). While none of our selected genes (*LAMA2*, *LSP1*, *MAD2L2*, *FBLN2* & *CXCL12*) appear to be associated with OS or prognosis of thyroid cancer in that study, events such as ES, AP and AT are concordant with those identified by *IsoformSwitchAnalyzeR()* in our HCC dataset. This suggests that while genes expressed in HCC are not previously implicated in thyroid cancer prognosis, the underlying splicing mechanisms may be shared among thyroid malignancy subtypes.

Despite being the first study to characterize functional consequences of isoform switching in HCC, there are several limitations of the study that should be acknowledged. This is reanalysis of a secondary dataset from NCBI, hence causal relationship between isoform switching and HCC pathogenesis cannot be deciphered. The analysis has been performed on patient cohort of 16 pairs. While this cohort size is adequate for statistical analysis, the isoform switches should be validated on large HCC patient’s cohort. Analysis was restricted to annotated sequences from Gencode v44 and novel transcripts relevant to HCC were ignored. The isoform switches and their functional consequences require experimental validation through in vitro techniques such as reverse transcription polymerase chain reaction (RT-PCR) and western blotting. Moreover, long read sequencing analysis is preferred for isoform switch analysis, so the identified switches in this study can be further validated through long read sequencing. Additionally, future studies should investigate whether the isoform switches identified here correlate with clinical outcomes including overall survival, recurrence and tumor stage, particularly as larger HCC-specific datasets become available. Despite the limitations, this study represents the first functional characterization of isoform switches in HCC and provides a foundation for future validation and experimentations.

## Supporting information

Combined Supplemental Tables

## Dataset and Code Availability

### Dataset availability statement

The original dataset and raw FASTQ files can be found at NCBI GEO with accession ID GSE228870.

### Additional Information

The complete script for Salmon quantification and isoform switch analysis can be found at the GitHub repository (https://github.com/razmia02/Isoform_Switch_Analysis).

## Author Contribution Statement

RB conceptualized the study, performed the bioinformatics analysis, compiled the datasets, and wrote the main manuscript text. RP & AA supervised the project, provided critical guidance on data interpretation, and revised the manuscript. Both authors reviewed and approved the final version of the manuscript.

## Supporting Information

### Supplementary Tables

**Supplementary Table S1.** Sample and metadata information.

**Supplementary Table S2.** Samples dropped due to low mapping rate.

**Supplementary Table S3.** QC metrics for all samples.

**Supplementary Table S4.** Switch consequence summary statistics.

**Supplementary Table S5.** Top 20 genes showing significant isoform switches with functional consequences.

**Supplementary Table S6.** Alternative splicing events summary statistics across isoforms used more and less in HCC.

**Supplementary Table S7.** Splicing enrichment results showing ATTS as statistically significant splicing mechanism.

**Supplementary Table S8.** Complete pathway enrichment analysis results from 335 genes.

## Acknowledgments

The authors would like to thank the researchers and primary investigators who generated and deposited the publicly available dataset used in this study (NCBI GEO Accession: GSE228870). We are grateful for their commitment to open science, which made this work possible.

## Competing Interest Statement

The authors have declared no competing interest.

## Notes

https://www.ncbi.nlm.nih.gov/geo/query/acc.cgi?acc=GSE228870

